# Consequences of the Nyquist-Shannon sampling criterion in Mesoscopic Multiphoton Microscopy to avail full-field sub-micron resolution resolvability

**DOI:** 10.1101/2021.01.31.429063

**Authors:** Bhaskar Jyoti Borah, Jye-Chang Lee, Han-Hsiung Chi, Yang-Ting Hsiao, Chen-Tung Yen, Chi-Kuang Sun

## Abstract

With a limited effective voxel rate, to date, each laser-scanning mesoscopic multiphoton microscope (MPM), despite securing an ultra-large field of view (FOV) and an ultra-high optical resolution simultaneously, experiences a *fundamental issue with digitization; i*.*e*., inability to satisfy the Nyquist-Shannon sampling criterion to resolve the optics-limited sub-micron resolution over the whole FOV. Such a system either neglects the criterion degrading the digital resolution to twice the pixel size, or significantly reduces the imaging area and/or the imaging speed to respect the digitization. Here we introduce a Nyquist figure of merit parameter to assess this issue, further to comprehend a maximum aliasing-free FOV and a cross-over excitation wavelength for a laser scanning MPM system. Based on our findings we demonstrate an ultra-high voxel rate acquisition in a custom-built *mesoscopic MPM system* to exceed the Nyquist-rate for a >3800 FOV-resolution ratio while not compromising the imaging speed as well as the photon-budget.

## 1. Introduction

Compared to single-photon and camera-based imaging systems, with a better penetration capability due to the use of near-infrared (NIR) excitation spectrum and excitation localization of nonlinear optical absorption, a laser scanning multiphoton microscope (MPM) becomes a promising candidate for deep intact tissue imaging while maintaining high enough 3D-resolution [1-10]. To accomplish high-speed imaging of a mesoscale volumetric tissue-sample, an extended FOV is a basic requirement. However, the FOV of a conventional MPM system is limited to <1 mm^2^ while preserving a sub-micron resolution. Recent reports from several researchers (Table 1) have addressed various design challenges to extend the FOV of a laser scanning MPM, and demonstrated their ultra-large FOVs up-to several square millimeters. In addition, by means of moderate- or high-numerical aperture (NA) objective lenses they preserved high optical resolutions as well. Even though an extended FOV of more than 1 mm^2^ and a sub-micron optical resolution are simultaneously achieved by means of a high-NA low-magnification objective lens, one must perform a proper digitization to retrieve the sub-micron structures. While doing so, to not sacrifice the imaging speed, a high voxel rate acquisition is essential to satisfy the Nyquist-Shannon sampling criterion [11-12], which requires *the digitized size of each sampling pixel to be at least half of the smallest resolvable spacing in order to prevent the phenomenon of aliasing, which essentially converts the optics-limited high spatial frequencies of the objects into low spatial frequencies in the final image by the Moiré effect* [13-14].

**Table 1.** A comparison of a few of the state-of-the-art large-FOV high-lateral-resolution MPM systems.

| Literature | NA of the objective | Axial resolution (optical) (μm) | Lateral resolution (optical) (μm) | Reported extended FOV (approx.) | FOV/resoluti on ratio | Reported fast-axis pixel number | NFOM (experi mental) |
| --- | --- | --- | --- | --- | --- | --- | --- |
| Tsai et al. [15] | 0.28 | 14 | 1 | 8 mm×10 mm | 8000 | 2000 | 0.125 |
| Bumstead et al. [16] | 0.22 | 28 | 1.7 | 4.95 mm×4.95 mm | 2912 | 1000 | 0.172 |
| Balu et al. [19] | 1.05 | 3.3 | 0.5 | 0.8 mm×0.8 mm | 1600 | 1600 | 0.5 |
| Stirman et al. [18] | 0.43 | 12.1 | 1.2 | 3.5 mm (width) | 2917 | 2048 | 0.351 |
| Terada et al. [17] | 0.6 | 9.96 | 1.26 | 1.2 mm×3.5 mm | 952 | 512 | 0.269 |
| This report | 0.95 | 2 @1070 nm | 0.48 @1070 nm | 1.6 mm×1.6 mm | >3000 | 8800 | ≥1.0 |
|  |  |  | 0.41 @919 nm | 1.6 mm×1.6 mm | >3800 |  |  |
|  |  |  | 0.37 @824 nm | 1.42 mm×1.42 mm | >3800 |  |  |
| **Note:** The NFOMs in this table [15-19] were calculated considering reported optical resolutions, following NFOM = 0.5 r<sub>opt</sub> P/ FOV; where, r<sub>opt</sub>, P, and FOV are reported lateral optical resolution, fast-axis pixel number, and fast-axis FOV, respectively; and P/ FOV is the inverse of the fast-axis pixel size. |  |  |  |  |  |  |  |

It is well-known that a high-NA objective lens is essential for high spatial resolution that can be further enhanced by lowering the excitation wavelength. Nevertheless, in case of digital imaging system the effective resolution no longer remains a function of only the NA and excitation wavelength, rather it highly depends on the capability of the respective system to perform a proper digitization to retrieve the finest optically-resolvable structures. In other words, failure to fulfill the basic Nyquist-Shannon criterion of digitization will end up a poor effective resolution irrespective of the high-NA and/or lower excitation wavelength being employed. In case of an extended FOV while maintaining a high lateral resolution, the requirement for fulfilling the Nyquist-Shannon criterion gets much tougher particularly when not sacrificing the imaging speed. This situation becomes more serious with a pulsed laser source for efficient nonlinear excitation, where the effective voxel rate is limited by the laser repetition rate, as each voxel must correspond to at least one optical pulse. As evident from the prior literatures [15-20], even though ultra-large FOVs were reported preserving ultra-high optical resolutions, as limited by inadequate effective voxel rates, none of the MPM systems could satisfy the Nyquist-Shannon sampling criterion to retrieve the best optical resolution across the ultra-large FOV. Consequently, the effective resolution of such a system degrades to two-times the voxel size, which indeed makes such an ultra-large FOV and ultra-high resolution not achievable at the same time in the final image, and enforces one to reduce the imaging speed and/or the imaging area for high resolution retrieval. To date, for a laser scanning MPM system, the realization of the Nyquist-Shannon sampling theorem to correlate its direct consequences over the maximum allowed FOV of such a system is yet unexplored. Especially for an MPM with an ultra-high FOV/resolution ratio, the issue of aliased digitization indeed persists as a barrier to unlock its ultimate potential and to enable an ultra-large aliasing-free FOV with a sub-micron effective resolution while preserving a high-speed raster-scanning.

In this paper, based on the Nyquist-Shannon sampling theorem we first formulate the minimum required repetition rate of a pulsed laser source to fulfill the Nyquist-Shannon criterion for a given laser scanning MPM system with a specific FOV. To characterize such an MPM system in terms of its reliable digitization capability, we introduce a Nyquist figure-of-merit (NFOM) parameter which indicates whether or not the system is capable of retrieving the best optical resolution. For the digitization to be aliasing-free, the value of NFOM must be greater than or at least equal to one. Taking NFOM into account, we then derive the maximum allowed FOV for a given laser-repetition-rate, fast-axis scanner frequency, excitation wavelength, and objective’s NA. Beyond this theoretical limit, the FOV will get aliased and the effective resolution will tend to degrade regardless of its superior optical design. For an MPM with an optimized optical FOV design, we further study the cross-over excitation wavelength, below which the FOV gets constrained by the respective NFOM and becomes wavelength dependent. Based on our derivation, we justify that in order to maximize the FOV while neither compromising the resolution nor the imaging speed, the key solution is to enable an ultra-high voxel-sampling rate by means of a high-repetition-rate pulsed laser source, where a one-pulse-per-voxel synchronized acquisition [21-22] is assumed for the optimum case. Our derivation further remarks that for a laser scanning MPM with a high-NA objective lens but employing a low-repetition-rate pulsed laser, it is not feasible to achieve a large millimeter-scale FOV without getting aliased unless the raster-scanning speed is greatly slowed down.

To validate our derivation experimentally, a design-optimized mesoscopic MPM system was custom-built (refer to the supplemental information) to yield an optics-limited FOV of up-to 1.6×1.6 mm^2^ while preserving a sub-micron lateral resolution with a 0.95 NA objective lens. By implementing a regular 70 MHz femtosecond laser following our design guideline, we successfully achieved aliasing-free MPM imaging, i.e., NFOM≥1, with an FOV/resolution ratio of more than 3000. To validate the optical-zoom-free sub-micron resolution retrievability, we performed two-photon imaging of biological tissue samples with fine enough structures, and reliably retrieved them without shrinking down the 1.6×1.6 mm^2^ imaging area if the excitation wavelength is longer than 912 nm. Our experimental study further confirms the Nyquist-Shannon sampling criterion in an MPM system, as well as the derived cross-over excitation wavelength. By shifting the excitation wavelength down to around 919 nm which is close to this cross-over wavelength, a resolution-optimized mesoscopic MPM system was also demonstrated with the 1.6×1.6 mm^2^ FOV, while maintaining NFOM=1 and ∼400 nm lateral resolution with a maximized FOV/resolution ratio of more than 3800. Whereas, our study further confirmed that with an excitation shorter than this cross-over wavelength, we had to shrink down the FOV to rescue an NFOM of at least 1, that is, the aliasing-free FOV in this regime is wavelength-dependent and is limited by Nyquist-Shannon criterion rather than the optical system design. Our study and demonstration of the mesoscopic MPM with an ultra-high FOV/resolution ratio can serve as a guideline to eliminate the final MPM barrier of aliased digitization and to enable deep-tissue volumetric MPM imaging with simultaneously an ultra-large aliasing-free FOV and a sub-micron digital resolution without compromising the imaging speed.

## 2. Results

To properly digitize the smallest optically resolvable spacing in a digital imaging system, the pixel size must be at least half of this smallest resolvable spacing, called the Nyquist-Shannon criterion [13-14], which must be satisfied to avoid the phenomenon of aliasing. For an objective lens with a specific value of NA, its smallest resolvable lateral spacing, i.e., the lateral resolution can be estimated by the full width half maximum (FWHM) of the lateral cross section obtained by imaging a small enough structure. Theoretically, for a high-NA (>0.7) objective lens, the lateral resolution (FWHM) for multiphoton fluorescence can be described as [23-24]-

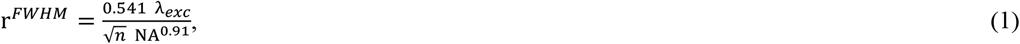

where λ_*exc*_ and *n* stand for excitation wavelength and order of the multiphoton process, respectively. In a pulsed-laser based point scanning MPM system, fulfilling/exceeding the Nyquist-Shannon criterion for an aliasing-free maximum fast-axis field of view of FOV_*max*_ with a specific lateral resolution of r^*FWHM*^ requires the repetition rate (R) of the pulsed laser source to be

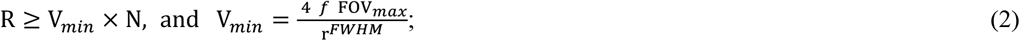

where, V_*min*_is the minimum voxel sampling rate required to fulfill the Nyquist-Shannon criterion, N (≥ 1) is an integer signifying number of optical pulse(s) per voxel, and *f* is the fast-axis frequency for either a resonant or a galvanometer-based scanner. Using Equation (1), V_*min*_can be redefined for an MPM system with a high-NA (>0.7) objective lens as

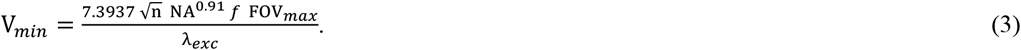

As V_*min*_is directly proportional to *f* and FOV_*max*_, and inversely proportional to λ_*exc*_, for a given value of V_*min*_, to extend the aliasing-free FOV while imaging with a high-NA objective lens, one must either decrease the fast-axis scanning frequency sacrificing the imaging speed, and/or increase the excitation wavelength sacrificing the resolution. Therefore, the only way left to neither compromise speed nor resolution is to enhance the voxel sampling rate sufficiently. For the optimized condition, following N = 1 in Equation (2) inequality, the lowest required repetition rate of the laser would thus be equal to the minimum voxel rate, i.e., R_*min*_ = V_*min*_. Thus, for a given MPM system, the required value of R_*min*_ can be estimated from Equation (3) based on desired FOV_*max*_ and choices of NA, *f*, and *λ*_*exc*_. For instance, if one employs an 8 kHz fast-axis scanner and a 1.0 NA objective lens for two-photon imaging at 920 nm excitation, for achieving a 2 mm wide fast-axis FOV, the minimum required voxel rate will be ∼181.86 M/s, and hence the laser repetition rate must be at least ∼181.86 MHz. This requirement can be of course relaxed to ∼90.9 MHz with a 4 kHz fast-axis scanner.

Considering Equations (2) and (3), for a straightforward assessment of the optical-resolution retrieving capability of a laser scanning MPM system, we formulate a Nyquist figure of merit (NFOM) parameter as follows

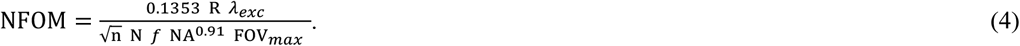

For the digitization to be aliasing-free, NFOM must be greater than or at least equal to 1. For a given *λ*_*exc*_, an attempt to enhance the FOV_*max*_ with a high-*f* and/or a high-NA configuration will tend to degrade the NFOM to be less than 1, which must be rescued by means of a high-R laser. Incorporating an even higher-NA objective lens, and/or lower *λ*_*exc*_ will demand an even higher value of R to maintain a specific FOV_*max*_. For instance, a ∼1.4 times higher value of R will be required when NA is enhanced from 1.0 to 1.45; likewise, lowering *λ*_*exc*_ from 1070 nm to 760 nm, will require a ∼1.4 times higher R, while the remaining parameters kept unchanged in each case.

For the optimized case with NFOM = 1 and N = 1, the aliasing-free FOV_*max*_ for a specified set of laser, fast-axis scanner, and objective lens can be thus estimated as

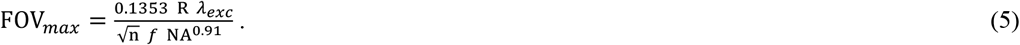

Figure 1*A* plots the maximum aliasing-free field of view, FOV_*max*_ from Equation (5) as a function of R for a fixed value of NA = 0.95 and N = 1. As depicted by the data points, to achieve an aliasing-free FOV_*max*_ of 1 mm to be operated up-to a minimum *λ*_*exc*_ of 920 nm, the laser repetition rate must be at least 43.4 MHz for a 4 kHz scanner, and at least 86.8 MHz for an 8 kHz scanner. This essentially justifies that, for a high-NA objective lens, employed with a high-frequency fast-axis scanner, the only way to extend the aliasing-free FOV_max_ beyond ∼1 mm is to opt for a pulsed laser source with high enough repetition rate (R) to enable an ultra-high effective voxel rate (V).

**Figure 1.**
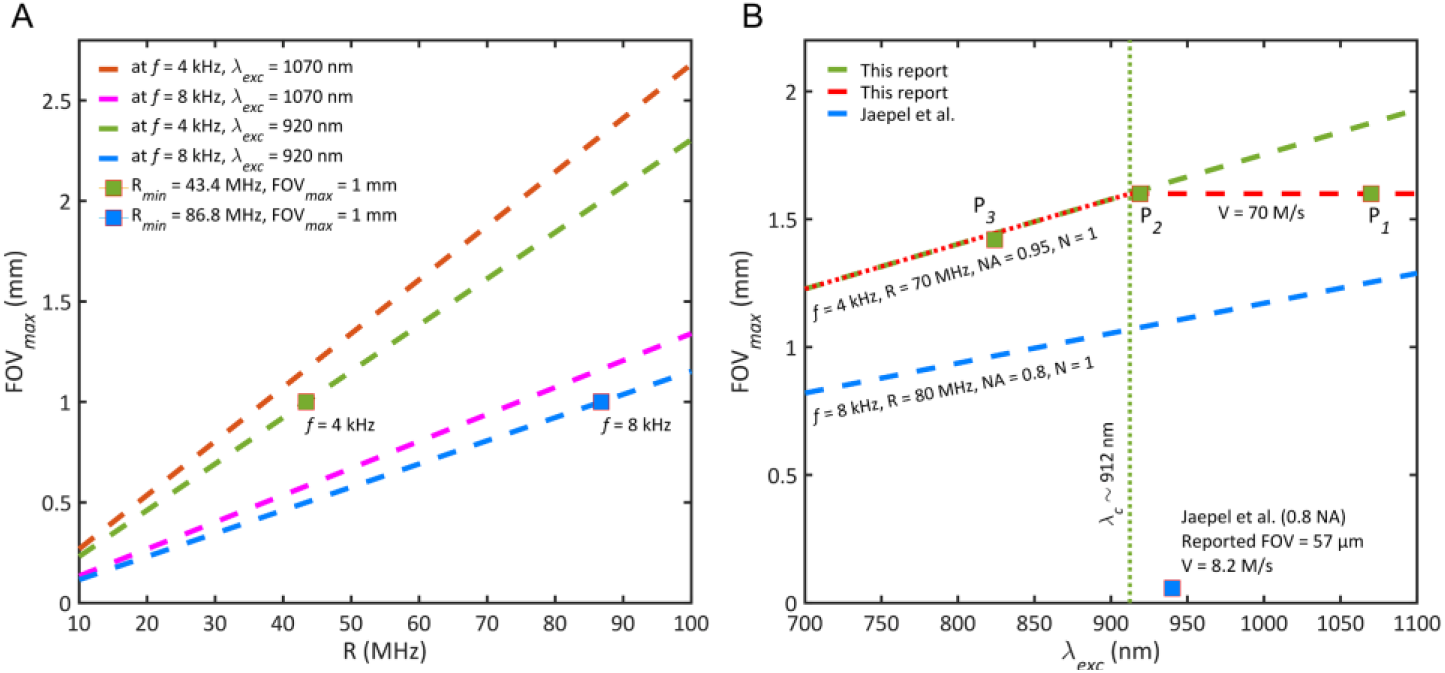
Plots of FOV_max_ for a laser scanning MPM system. (A) FOV_max_ with respect to R at NA = 0.95 & N = 1. For FOV_max_ = 1 mm, the green & blue colored solid squares correspond to R = 43.4 MHz at f = 4 KHz & R = 86.8 MHz at f = 8 KHz, respectively. (B) FOV_max_ with respect to λ_exc_, where the green-dashed line represents FOV_max_ of this report at NA = 0.95, N = 1, and f = 4 KHz; green-dotted line marks the cross-over excitation wavelength, i.e., λ_c_ = 912 nm; red-dashed and red-dotted lines depict the optics-limited and NFOM-constrained FOVs, respectively. P_1_, P_2_, and P_3_ are three imaging conditions each with 70 M/s effective voxel rate, at excitation wavelengths of 1070 nm (> λ_c_) with NFOM > 1 for 1.6 mm-wide optics-limited FOV, at 919 nm (~λ_c_) with NFOM = 1 for the same 1.6 mm-wide but resolution-optimized FOV, and at 824 nm (< λ_c_) with NFOM = 1 for an NFOM-constrained 1.42 mm-wide FOV, respectively. Blue-dashed line represents theoretical FOV_max_ limitation for the experimental condition of a prior report by Jaepel et al. [25] with an 80 MHz pulsed laser. The blue solid square marks the previous experimental FOV [25], which is much lower than the theoretical limitation derived in this paper.

Based on Equation (5), Figure 1*B* plots the FOV_*max*_ as a function of *λ*_*exc*_ for two high-NA MPM system settings each with NFOM≥1, where the blue-dashed line corresponds to the experimental parameters of a prior report by Jaepel et al. [25]. In this case, although an 80 MHz pulsed laser was employed for excitation, the previously reported aliasing-free FOV is >19 times lower than our idealized theoretical value FOV_*max*_ with N=1 (green-dashed line). Part of the discrepancy can be attributed to the low voxel sampling rate of 8.2 M/s. On the other hand, our report studied the case of a femtosecond laser a with 70MHz repetition rate, which is on the same order or slightly lower than most commercial femtosecond lasers. As shown in the green-dashed line, which represents FOV_*max*_ with NA = 0.95, N = 1, and *f* = 4 KHz, with a maximized effective voxel rate of 70 M/s this report predicts the possibility to achieve an aliasing-free FOV_*max*_ much greater than 1 mm across the excitation wavelengths between 700-1100 nm while digitally preserving the diffraction-limited nonlinear optical resolution. Our calculation also justifies the impact of true voxel rate (V) and hence the laser repetition rate (R) for a large-FOV aliasing-free imaging.

For a conventional laser scanning system [26] with an optics-limited field of view of FOV_*OL*_, we can redefine the minimum effective voxel rate, and hence the minimum laser-repetition-rate required for a given laser scanning MPM system to be aliasing-free. Following Equation (3), and considering FOV_*max*_ = FOV_*OL*_, we obtain

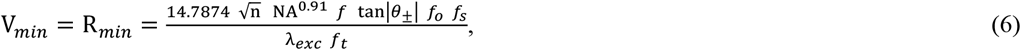

where, *θ*_±_ is the fast-axis scan-angle by the scanning mirror with respect to the optical axis, *f*_*o*_, *f*_*s*_, and *f*_*t*_ are the effective focal lengths (EFLs) of the objective lens, scan lens, and tube lens, respectively.

From Equation (5), for a laser scanning MPM system with FOV_*OL*_ ≤ FOV_*max*_, a cross-over excitation wavelength *λ*_*c*_ can be further estimated as follows. When *λ*_*exc*_ is longer than *λ*_*c*_, the FOV of the system remains limited by the relevant optics. However, as *λ*_*exc*_ becomes shorter than *λ*_*c*_, the FOV is rather constrained by the NFOM of the respective system. For an optics-limited resolution-optimized-FOV, *λ*_*exc*_~*λ*_*c*_ is expected with

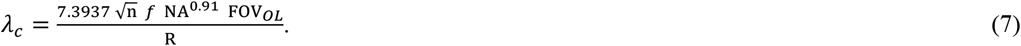

To validate our theory and hypothesis, we constructed a mesoscopic MPM system utilizing off-the-shelf optomechanical components. Following Equation (6), we chose *f*_*o*_, *f*_*s*_, and *f*_*t*_ of 9 mm, 110 mm, and 166.7 mm, so that *θ*_±_~ ± 7.7^0^ can provide up-to an FOV_*OL*_ of ∼1.6 mm. Combining a high (0.95) NA objective lens with 9 mm EFL preserved not only a high spatial resolution, but also resulted in a large FOV/resolution ratio greater than 3000. For two-photon imaging we further chose *λ*_*exc*_ = 1070 *nm* and *f* = 4 *KHz*, thus obtaining V_*min*_ = R_*min*_ = 59.7 MHz, which is lower than most commercially available femtosecond oscillators and is easily achievable. For demonstration, a femtosecond laser source with R = 70 MHz and *λ*_*exc*_ ≈ 1070 *nm* was first chosen. Following Equation (2), the value of N, being an integer, is not allowed to be more than 1 in this case, enforcing one-pulse-per-voxel acquisition [21-22]. Therefore, following Equation (5), we obtain FOV_*max*_ = 1.87 *mm* (> FOV_*OL*_) for N = 1 case. From Equation (4), NFOM = 1.17 > 1, thus, the optics-limited FOV is guaranteed to be aliasing-free. In this case, from Equation (7), the cross-over excitation wavelength, *λ*_*c*_ was estimated to be around 912 nm.

To experimentally validate the fulfillment of the Nyquist-Shannon criterion, we performed two-photon imaging of a coronal section from the medulla of a Nav1.8-tdTomato mouse across a volume size of ∼1.6×1.6×0.4 mm^3^ preserving a voxel size of 0.182×0.182×0.3 μm^3^. With *λ*_*exc*_ ≈ 1070 *nm* > *λ*_*c*_, the FOV was optics-limited in this case. Figure 2*A* depicts the stitch-free 3D rendered volume (Amira; Mercury Computer Systems) in an inclined view. The 3D region-of-interest (ROI) R1 is cropped and enlarged in Figure 2*B*, and its top view is presented in Figure 2*C*. Red, green and cyan colored axes represent X, Y and Z axes, respectively. Figure 2*D* shows a 2D representation of the acquired volume with color-coded depth information, processed using ImageJ software (NIH, Bethesda, MD) and OpenCV (an open-source computer vision library). No stitching/ mosaicking was applied. Figure 2*E* shows a 3× digitally zoomed ROI R2 with 2933×2933 pixels maintaining a ∼182 nm pixel size, presenting the color-coded fine fibers. With a 26× digital zoom to the original image, the ROIs R3 and R4 marked in Figure 2*E* are enlarged in Figures 2 *F* and *G*, respectively, each with a pixel number of 333×333, with 182 nm pixel size. The red arrows in Figures 2 *F* and *G* mark two sub-micron fibers. Figures 2 *H* and *I* respectively show the intensity profiles across the fibers in Figures 2 *F* and *G*, with FWHMs of around 475 nm and 470 nm, respectively. Figure 2*J* depicts a Z-projected ∼9 μm-thick section from the same acquired data with pixel size of 182 nm. An 8.5× digital zoom was performed to the ROI R5 (1161×912 pixels) and its enlarged view is shown in the same Figure 2*J*. Another ROI R6 with two adjacent submicron fibers is selected within ROI R5, with a 54× digital zoom to the original image. Figures 2 *K-N* respectively depict the enlarged views of the ROI R6 with (*K*) Nyquist-exceeded (pixel size = 182 nm), (*L*) 2x undersampled (pixel size 364 nm), (*M*) 3x undersampled (pixel size 545 nm), and (*N*) 4x undersampled (pixel size 727 nm) cases (scale bar = 5 μm), with pixel numbers of 226×117, 112×58, 75×38, and 56×29, respectively. ROI R6 in *J* is vertically oriented in *K-N*. Figure 2*O* plots the intensity profile along the yellow-dashed line marked in Figure 2*K* for the Nyquist exceeded condition, and resolves the separation between the two adjacent sub-micron fibers which are ∼550 nm apart. On the contrary, similarly plotted intensity profiles in Figures 2 *P-R*, for the undersampled cases, essentially detect these two adjacent fibers as a single fiber, as the increased pixel size does not meet the Nyquist-Shannon criterion.

**Figure 2.**
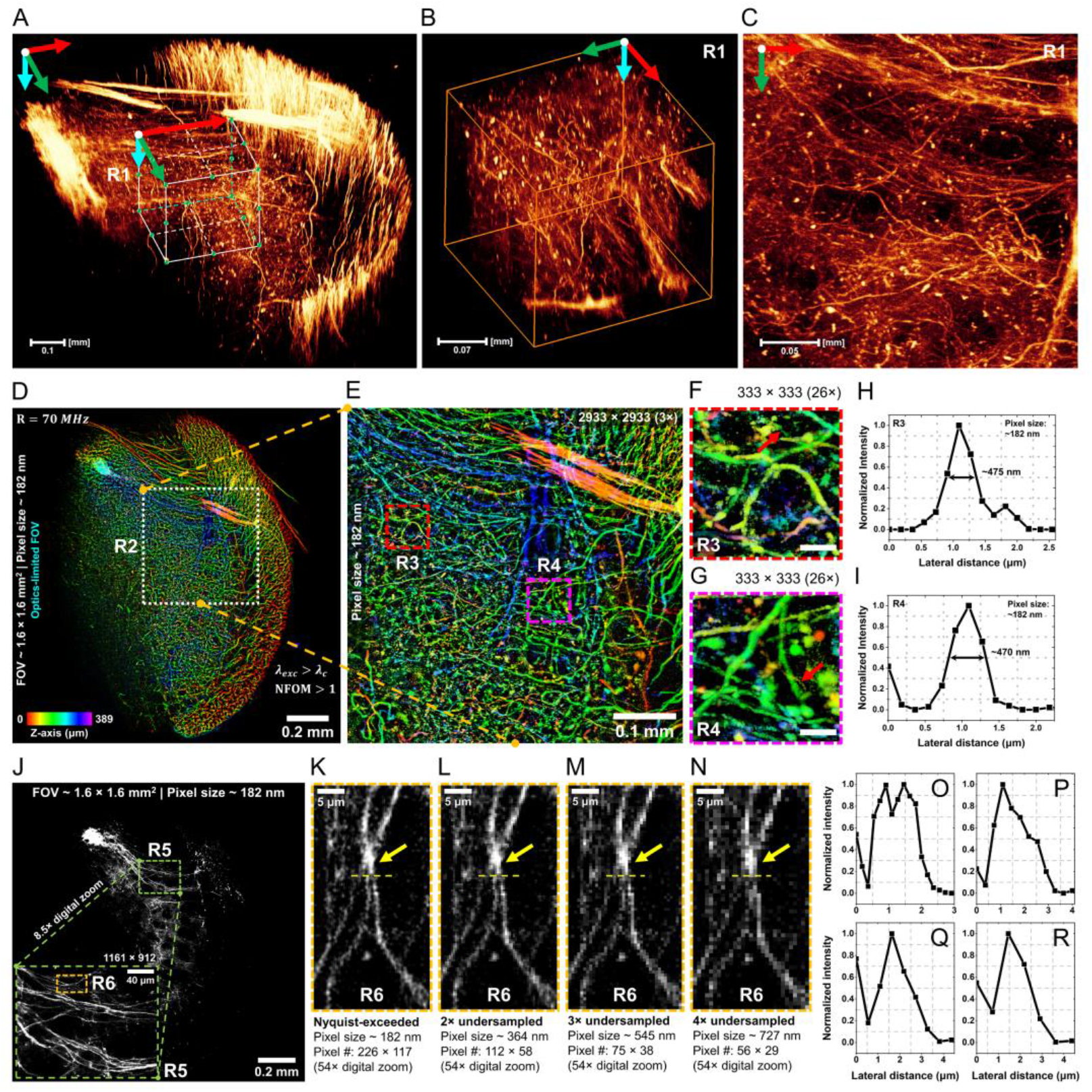
Demonstration of stitch-free large-FOV two-photon imaging with Nyquist-exceeded sampling. A coronal section from medulla of a Nav1.8-tdTomato mouse was scanned across 1.6×1.6×0.4 mm^3^ volume with a voxel size of 0.182×0.182×0.3 μm^3^. (A) 3D rendered volume (using Amira software) in inclined view; (B) enlarged 3D cropped ROI R1 marked in A, providing lateral & axial views; (C) top view of the ROI R1. Red, green & cyan colored axes represent X, Y & Z axes, respectively. Scale bars for A-C: 0.15 mm, 0.07 mm, and 0.05 mm, respectively. (D) 2D representation of the acquired volume with Z-projection (color coded Z-axis), scale bar = 0.2 mm. 3× digitally zoomed ROI R2 (marked in D), scale bar = 0.1 mm, showing color-coded fine fibers; & (G) 26× digitally zoomed ROIs R3 & R4 in E; red arrows mark sub-micron structures, scale bar = 15 μm. (H) & (I) Intensity profiles across the fine fibers marked in F & G, respectively; revealing FWHMs of 475 nm for H, and 470 nm for I. (J) A Z-projected 9 μm-thick section from the same acquired data, scale bar = 0.2 mm. (K-N) enlarged vertically oriented views of the ROI R6 marked inside ROI R5 in J, obtained with (K) Nyquist-exceeded (pixel size 182 nm), (L) 2x undersampled (pixel size 364 nm), (M) 3x undersampled (pixel size 545 nm), and (N) 4x undersampled (pixel size 727 nm) cases, respectively (scale bar = 5 μm), with pixel numbers of 226×117, 112×58, 75×38, and 56×29, respectively; (O-R) Intensity profiles along the yellow-dashed lines in K-N, respectively. (O) resolves the separation between two adjacent sub-micron fibers. (P-R) essentially detect the two adjacent fibers resolved in O as a single fiber.

In Figure 1*B*, the green-dashed line corresponds to the FOV_*max*_ of our custom-built MPM system plotted with respect to the excitation wavelength. For the 70 MHz laser, the cross-over excitation wavelength *λ*_*c*_ in our case is around 912 nm as represented by the green-dotted line in Figure 1*B*. In Figure 2 with an NFOM>1, we have demonstrated the Nyquist-exceeded imaging capability of the system at *λ*_*exc*_ of around 1070 nm as indicated by point-P_*1*_ in Figure 1*B*. At this condition with *λ*_*exc*_ > *λ*_*c*_, the 1.6 mm-wide FOV was limited by optics and was lower than the FOV_*max*_ limit. The red-dashed line in Figure 1*B* represents our optics-limited FOV.

To realize not just the optics-limited but also resolution-optimized FOV, the excitation wavelength was lowered to around 919 nm (point-P_*2*_ in Figure 1*B*) approaching the cross-over wavelength (*λ*_*c*_) of 912 nm. We performed two-photon imaging of a sagittal section from the hippocampus of a thy1-GFP mouse. The Z-projected view of 8 imaging slices at 0.8 μm Z-step is depicted in Figure 3*A* with a scale bar of 0.2 mm. An NFOM=1 for the optics-limited 1.6 mm-wide FOV was maintained with the same pixel size of 182 nm. To investigate the high-resolution resolvability, we first mark an ROI R1 in Figure 3*A* with 512×512 pixels which is enlarged in Figure 3*B* with a scale bar of 20 μm. Another ROI R2 with 64×64 pixels is again marked inside ROI R1 in Figure 3*B*. Figure 3*C* shows the enlarged view of R2 with a scale-bar of 2 μm which resolves fine sub-micron structures by means of the 138× digital zoom to the original image. An intensity profile obtained across the fiber marked by the white arrow in Figure 3*C* is plotted in Figure 3*D* with an FWHM of 405 nm. Thereby, an FOV/resolution ratio of more than 3800 was enabled, not to mention without optical zooming.

**Figure 3.**
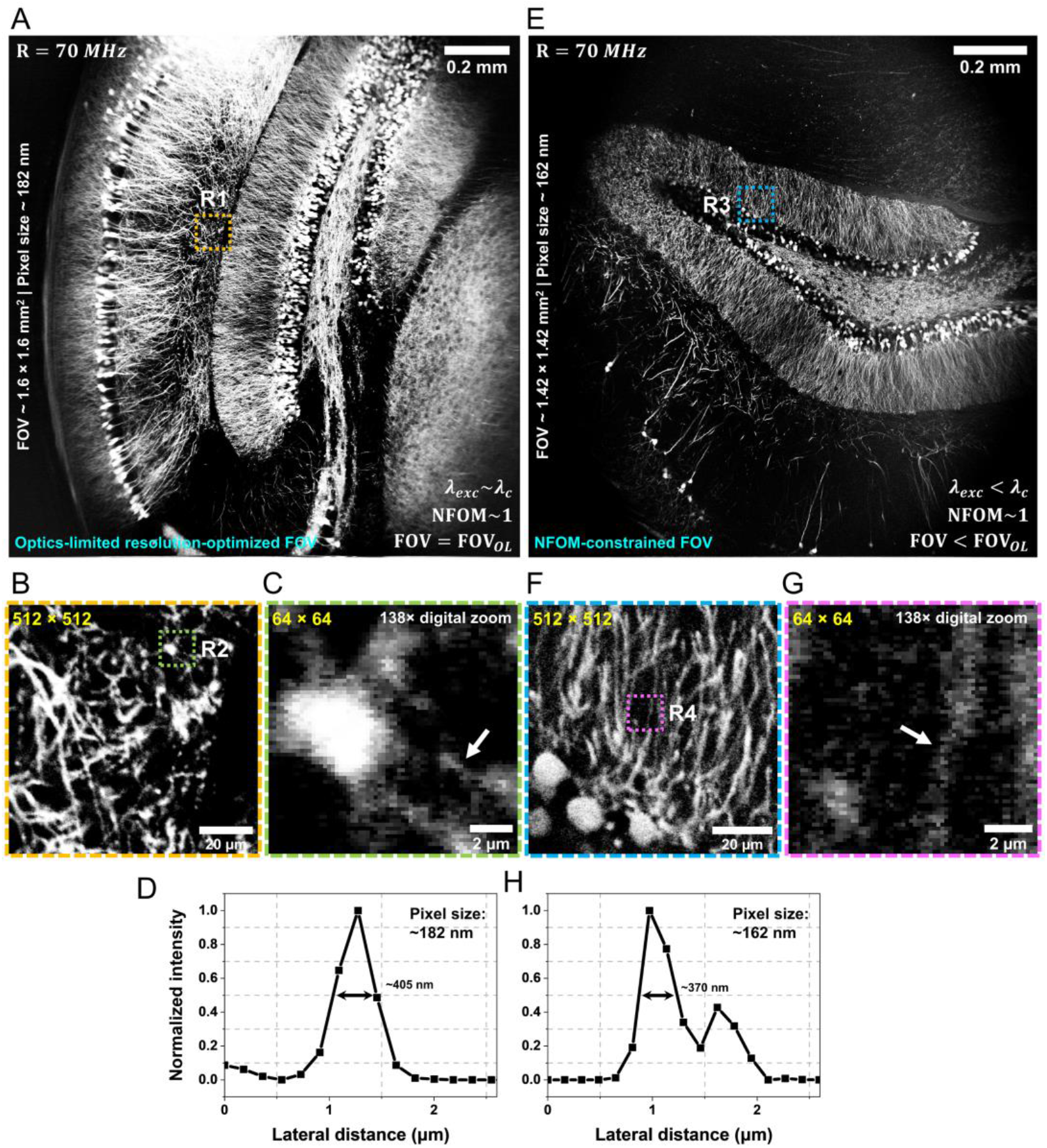
Two-photon imaging near & below the cross-over excitation wavelength to demonstrate an optics-limited resolution-optimized FOV & an NFOM-constrained FOV, respectively. (A) Optics limited 1.6 mm-wide FOV at λ_exc_ = 919 nm ~ λ_c_ & NFOM = 1; Z-projected view of 8 imaging slices at Z-step = 0.8 μm, pixel size = 182 nm, scale bar = 0.2 mm; sample: a hippocampal sagittal section from a thy1-GFP mouse. (B) & (C) Enlarged ROIs R1 (512×512 pixels) in A & R2 (64×64 pixels) in B, respectively. (D) Intensity profile across the fiber marked by the white arrow in C with an FWHM of 405 nm. (A-D) justifies the best resolution for the optics-limited FOV with an FOV/resolution ratio of more than 3800 at the optimized condition with NFOM=1 & λ_exc_~ λ_c_. (E) NFOM-constrained 1.42 mm-wide FOV at λ_exc_ = 824 nm < λ_c_; Z-projected view of 8 imaging slices at Z-step = 0.8 μm, pixel size = 162 nm, scale bar = 0.2 mm; sample: a hippocampal coronal section from an Alexa Fluor 546 (anti-GFP immunohistochemistry labeling) mouse. (F) & (G) Enlarged ROIs R3 in E with 512×512 pixels & R4 in F with 64×64 pixels respectively. (H) Intensity profile across the white-arrow-marked fiber in G revealing an FWHM of 370 nm comparable to the respective r^FWHM^ of 330 nm. Scale bars in B & F are 20 μm, and C & G are 2 μm. Note: ROIs R2 & R4 in C & G, respectively, were obtained by performing 138× digital zoom to the corresponding full-FOV images.

To demonstrate the predicted effect of NFOM-constrained FOV as depicted by the red-dotted line in Figure 1*B*, we again reduced the excitation wavelength to around 824 nm which is lower than *λ*_*c*_. At this condition, the aliasing-free FOV is no longer limited by the optical performance, rather it becomes NFOM-constrained which in our case is around 1.45 mm. Remarkably, such NFOM-constrained reduction to the FOV is going to be more severe for an application requiring even shorter excitation wavelength. To demonstrate this scenario, we acquired two-photon images from a coronal section of an Alexa Fluor 546 stained (anti-GFP immunohistochemistry labeling) mouse hippocampal region at *λ*_*exc*_ of around 824 nm as represented by point-P_*3*_ in Figure 1*B*. Figure 3*E* with a scale bar of 0.2 mm depicts the Z-projected view of 8 imaging slices at 0.8 μm Z-step. Respecting the NFOM-constrain, the FOV was set to 1.42 mm with a reduced pixel size of 162 nm. The 512×512 pixeled ROI R3 in Figure 3*E* is enlarged in Figure 3*F* (scale bar = 20 μm) where another 64×64 pixeled ROI R4 is again marked. Figure 3*G* depicts the enlarged view of the ROI R4 with a 138× digital zoom to the full-FOV image. Figure 3*H* plots the intensity profile across the white-arrow-marked fine fiber in Figure 3*G*, revealing an FWHM of around 370 nm being comparable to the theoretical r^*FWHM*^ of 330 nm.

Notably, both *λ*_*exc*_~*λ*_*c*_ case with optics-limited resolution-optimized-FOV and *λ*_*exc*_ < *λ*_*c*_ case with NFOM-constrained FOV preserved an NFOM∼1, and thus 2 pixels per theoretical point spread function was ensured in each case. ImageJ and OpenCV was used for the analysis presented in Figure 3.

## 3. Discussion

The purpose of an extended FOV while simultaneously preserving a high spatial resolution is to enable one to resolve fine sub-micron structures throughout the FOV without optical zooming, and thereby to facilitate a faster imaging speed with minimal digital image stitching. However, for an MPM system with an ultra-high FOV/resolution ratio together with a high enough raster-scanning speed, fulfillment of the Nyquist-Shannon sampling criterion becomes challenging due to requirement of an ultra-high effective voxel rate. A comparison of a few of the state-of-the-art large-FOV high-lateral-resolution MPM systems [15-19] in terms of FOV-resolution-ratios and NFOMs is enlisted in Table 1. Remarkably, each prior report with an inadequate voxel rate encountered an NFOM<1, and thus a degraded effective resolution in each case. Sofroniew et al. [20] demonstrated a 5 mm-wide FOV by means of combining multiple 600 μm-wide strips, however, was limited by pixel size to meet the Nyquist-Shannon criterion for the 0.66 μm reported resolution. Tsai et al. [15] reported an ultra-high FOV-resolution-ratio of 8000 preserving a 1 μm lateral resolution by means of a low (≤0.3) NA objective lens, however encountered a poor NFOM of 0.125. A lower value of NA typically leads to a poor axial resolution. Besides, a poor NFOM induces substantial degradation to the effective lateral resolution as well. Therefore, a low-NA poor-NFOM system results in a severely poor effective 3D resolution.

To achieve NFOM≥1 for our custom built MPM, the minimum required effective voxel rate was estimated to be 59.7 MHz (Equation (3) or (6)). Therefore, with a 70 MHz pulsed laser, Equation (2) restricts us to follow N = 1, i.e., a one-pulse-per-voxel synchronized acquisition to meet the voxel rate requirement. Despite being a promising idea, to date, this has not been implicated in a laser-scanning MPM aiming to maximize the effective voxel rate so as to fulfil/exceed the Nyquist-Shannon criterion for a large FOV/resolution ratio. It is remarkable that, the prior literatures related to pulse-level synchronized-sampling targeted small-FOV MPM systems, and either employed a low-repetition-rate pulsed laser source [21-22], or lowered down the effective voxel rate [27-30], basically to improve the signal to noise ratio (SNR). A low-repetition-rate laser is usually employed when aiming to a high pulse energy to secure a high SNR. Furthermore, whether the sampling being pulse-synchronized or not, a reduction to the effective voxel rate by means of any interpolation and/or pixel-binning method essentially degrades the NFOM irrespective of a high-SNR and can induce irreversible resolution loss. *On the other hand, our idea contradicts both the prior trends, i*.*e*., *usage of a low repetition-rate laser source and/or reduction of the effective voxel rate especially when a non-aliased extended FOV is a concern*. Our idea of maximizing the aliasing-free FOV enforces a maximized effective voxel rate limited by the repetition rate of a high pulse-rate laser at N = 1 for the acceptable condition.

For a conventional MPM with N = 1, due to lower pixel dwell time, a reduced photon budget, and hence a poor SNR can be a cause of concern. However, for a mesoscopic MPM with a large FOV, this situation is different. With N = 1, the reduced photon budget can be compensated by a higher excitation power without damaging the sample; since, the average power is now covering much extended area, and thus the power density over a unit area will decrease and will allow higher average power after the objective lens. For instance, by extending the FOV from 0.4×0.4 mm^2^ to 1.6×1.6 mm^2^, 16 times average output power will be needed to maintain the same optical power density over the unit area. For N = 1, there will be only two consecutive pulses to be focused inside the point spread function, and thus even with a time interval on the order of the fluorescence lifetime, the risk of bleaching is minimized. Our study thus yields a conclusion that for an ideal system with NFOM = 1 and N = 1, a high enough repetition-rate-laser with reasonably increased excitation power will allow the photon budget issue to be improved without much compromising the imaging speed. We utilized an average power of ≤45 mW for the experiments presented in this paper. In addition to the excitation power, collection of at least 8 voxels per focal volume for a Nyquist-exceeded volumetric imaging further improves the SNR. (See *Materials and Methods, 1*.*4* for SNR analysis at different frame accumulation conditions.) With no frame accumulation, an SNR over 8 was achieved, which was further improved to over 20 with a frame accumulation of 3.

We stated the repetition rate (R) of the pulsed laser source to be the only enhanceable parameter for achieving NFOM≥1 while not compromising with the imaging speed and/or the effective resolution. For a low-repetition-rate pulsed laser, one must reduce the fast-axis scanner frequency significantly to make NFOM≥1. For instance, following Equation (5) for a two-photon process with R = 1 MHz, *λ*_*exc*_ = 1070 *nm*, NA = 0.95, and FOV_*max*_ = 1.6 *mm*, to achieve NFOM = 1, the maximum allowed fast-axis frequency (*f*) is 67 Hz only, taking 7.46 ms per fast-axis line. Hence, for a square FOV of 1.6×1.6 mm^2^, a total of 55.64 seconds per frame will be required to fulfill the Nyquist-Shannon criterion with a pixel number of 7459×7459 for an r^*FWHM*^ of 429 nm, leading to a severely poor imaging speed of 0.018 fps. Hence, to maintain a large-field high-resolution imaging at high-speed with a resonant fast-axis scanner, and to simultaneously maintain NFOM≥1, a high repetition-rate pulsed laser is the only key. The imaging speed can be further enhanced by employing an even faster fast-axis scanner which will however demand an even higher repetition-rate pulsed laser to maintain NFOM≥1. However, most of the fluorescent dyes have a fluorescence lifetime of 1-5 ns, and therefore the pixel dwell time for point scanning needs to be >5 ns, and hence the value of R prefers not to be much higher than 200 MHz [31].

Even though we are not able to build a system capable of providing an aberration-free and vignetting-free extended-FOV with a full-field diffraction-limited performance, our validation of the Nyquist-exceeding capability in the central area of the FOV is a proof of the Nyquist-exceeding capability across the entire FOV, since the actual digital pixel size at the edge area is the same or even smaller than that in the central area due to the slower scanner speed and the spatial resolution at the edge is either the same (for a future ideal system) or poorer than that in the central imaging area.

With our demonstrated mesoscope, no optical zooming was employed to obtain the high-magnification images presented in this study. Instead, all sub-micron structures were retrieved by digitally zooming into the respective full-FOV images. Each pixel size we specified was identical in both horizontal and vertical directions, hence, the Nyquist-Shannon criterion was satisfied in both X and Y axes identically. Additionally, we used the theoretical resolution of the system in Equation (1) to evaluate our NFOM, ensuring digitization of all regions of the extended FOV fulfills the Nyquist-Shannon criterion. In the axial direction, a step-size of ≤800 nm was required to retrieve the best axial resolution of the system, and to ensure a collection of at least 8 voxels per focal volume. The axial movement speed was limited by the electronic stage OSMS80-20ZF-0B (SIGMA KOKI, Tokyo, JAPAN), providing a maximum travel of up-to 1 mm/sec.

In summary, we correlated the Nyquist-Shannon sampling theorem in laser scanning multiphoton microscopy to realize its impact over the field of view and its direct relationship with the excitation wavelength, the laser-repetition-rate, and the fast-axis scanning frequency. We formulated a Nyquist figure-of-merit parameter to characterize a laser scanning MPM in terms its reliable digitization capability. We defined the maximum allowable FOV for such a system which must not be exceeded to prevent digital resolution loss. We defined a cross-over excitation wavelength, which must not be subceeded to prevent NFOM-constrained reduction to the optics-limited FOV.

Based on our derivation, we proposed the way to maximize an aliasing-free FOV yet not compromising with the imaging speed and/or the effective resolution by enabling a laser-repetition-rate-limited maximized effective voxel rate. Remarkably, our idea contradicts the usage of a lower-pulse-rate laser when a non-aliased extended FOV with a high enough lateral resolution and a fast-enough imaging speed become the concerns.

We applied our idea in a custom built MPM system with an FOV/resolution ratio of more than 3000 and successfully demonstrated its optical-zoom-free sub-micron resolution retrieving ability by performing two-photon imaging of fine enough structures. We showed two-photon imaging above, near and below the cross-over excitation wavelength, where in the first two cases the FOV remained optics-limited, whereas, in the last case FOV was rather constrained by the NFOM.

This paper will help one to realize the consequences of the Nyquist-Shannon sampling theorem in context of a laser scanning multiphoton microscope. The Nyquist-exceeded mesoscopic imaging at sub-micron effective resolution presented in this paper will enable one to point-scan a large-volumetric sample, for instance an intact whole mouse brain preserving an optics-limited resolution with a reduced requirement of digital image stitching, not to mention without employing an optical-zoom for high resolution retrieval. Apart from volumetric imaging, it further holds a tremendous potential for rapid giga-pixel imaging of large-area *ex-vivo* biopsy tissue-samples to enable a real-time visualization of fine sub-micron histopathological features without optical zooming.

## Acknowledgments

This project was supported by Ministry of Science and Technology (Taiwan) with financial grants MOST 107-2221-E-002-157-MY3 and MOST 107-2321-B-002-006. We thank Dr. Daniel Lin (SunJin Lab Co., Taiwan) for advice and support in sample preparation.

## Author Contributions

B. J. Borah designed, optimized and implemented the complete optical and electronic systems; developed the C++ based control and data acquisition software; further performed imaging and data analysis. J.-C. Lee, H.-H. Chi and C.-T. Yen were in charge for the preparation of the biological samples used in this study. Y.-T. Hsiao prepared the lower wavelength two-photon excitation source. C.-K. Sun initiated the concept and conducted the research. B. J. Borah and C.-K. Sun wrote the paper.

## Methods

### Preparation of the biological test samples

The animals were maintained in accordance with guidelines approved in the Codes for Experimental Use of Animals of the Council of Agriculture of Taiwan, based on the Animal Protection Law of Taiwan. All experimental protocols were approved by the Institutional Animal Care and Use Committee of National Taiwan University, Taipei, Taiwan. The transgenic Nav1.8-tdTomato male mice used in this study were 8-week-old. They were housed with a 12-hour light/12-hour dark cycle and fed ad libitum. For the sample preparation, mice were processed with passive CLARITY method [32]. The mice were anesthetized with overdose of sodium pentobarbital (100 mg/kg) and perfused transcardially with ice-cold phosphate buffered saline, and followed with hydrogel monomer (4% acrylamide, 2% bis-acrylamide, 4% paraformaldehyde and VA-044 initiator). The brains and sciatic nerves with dorsal root ganglion were dissected and incubated at 4° C for 2 days, and then polymerized at 37° C for 3 hours. After removing the extra hydrogel from the surface, the brain and nerve samples were washed on a rotating shaker with 4% sodium dodecyl sulfate (SDS) clearing solution at room temperature. The clearing solution was replaced each week and the tissue clearing status was monitored. The passive CLARITY clearing procedure was performed up-to 4 months for the whole-brain clarifying samples, while one week was enough for the sciatic nerve sample. After the passive CLARITY clearing procedure, the brain samples were washed in PBST (0.3% Triton X-100 in phosphate buffered saline) for 3 days to remove the SDS clearing solution. For the ∼500 µm medulla section, one brain was sliced by a vibratome. We used the RapiClear CS (refractive index 1.45; SunJin Lab Co., Taiwan) for refractive index matching overnight, and then embedded the medulla sample, whole-brain sample, and nerve sample within suitable spacers (iSpacer; SunJin Lab Co., Taiwan) for imaging. Additionally, for the experiments with GFP and Alexa Fluor 546, two thy1-GFP mice were first transcardially perfused with ice-cold phosphate buffered saline, and followed with 4% paraformaldehyde. The brains were dissected and post-fixed at 4^0^ C for 2 days. The 250 µm sections were prepared by a vibratome in sagittal or coronal orientation. One coronal section was stained with immunohistochemistry procedures: rabbit ant-GFP antibody (ThermoFisher, A-11122, 1:200), Goat-anti-Rabbit-Alexa 546 (ThermoFisher, A-11035, 1:400). The sections were processed with RapiClear 1.52 (SunJin Lab Co., Taiwan) for refractive index matching.

## References

1. Denk, W., Strickler, J. H., and Webb, W. W., Two-photon laser scanning fluorescence microscopy. Science 248, 73–76 (1990).

2. Helmchen, F., and Denk, W., Deep tissue two-photon microscopy. Nat. Methods 2, 932–940 (2005).

3. Theer, P., Hasan, M. T., and Denk, W., Two-photon imaging to a depth of 1000 µm in living brains by use of a Ti:Al2O3 regenerative amplifier. Opt. Lett. 28, 1022–1024 (2003).

4. Ntziachristos, V., Going deeper than microscopy: the optical imaging frontier in biology. Nat. Methods 7, 603–614 (2010).

5. Jacques, S. L., Optical properties of biological tissues: a review. Phys. Med. Biol. 58, 37–61 (2013).

6. Kobat, D., Durst, M. E., Nishimura, N., Wong, A. W., Schaffer, C. B., and Xu, C., Deep tissue multiphoton microscopy using longer wavelength excitation. Opt. Express 17, 13354–13364 (2009).

7. Horton, N. G., Wang, K., Kobat, D., Clark, C. G., Wise, F. W., Schaffer, C. B., and Xu, C., In vivo three-photon microscopy of subcortical structures within an intact mouse brain. Nature Photon 7, 205–209 (2013).

8. Horton, N. G., and Xu, C., Dispersion compensation in three-photon fluorescence microscopy at 1,700 nm. Biomed. Opt. Express 6, 1392–1397 (2015).

9. Rosenegger, D. G., Tran, C. H. T., LeDue, J., Zhou, N., and Gordon, G. R., A High Performance, Cost-Effective, Open-Source Microscope for Scanning Two-Photon Microscopy that Is Modular and Readily Adaptable. PLoS ONE 9, e110475 (2014).

10. Chakraborty, S., Lee, S. Y., Lee, J. C., Yen, C. T., and Sun, C. K., Saturated two-photon excitation fluorescence microscopy for the visualization of cerebral neural networks at millimeters deep depth. J. Biophotonics 12, e201800136 (2019).

11. Shannon, C. E., Communications in the presence of noise. Proc. IRE 37, 10–21, (1949).

12. Nyquist, H., Certain topics in telegraph transmission theory. Trans. AIEE 47, 617–644 (1928).

13. Pawley, J. B., Handbook of Biological Confocal Microscopy. Ed. (J. B. Pawley, New York: Springer), pp. 59–79 (2006).

14. Heintzmann, R., and Sheppard, C. J. R., The sampling limit in fluorescence microscopy. Micron 38, 145–149 (2007).

15. Tsai, P. S., Mateo, C., Field, J. J., Schaffer, C. B., Anderson, M. E., and Kleinfeld, D., Ultra-large field-of-view two-photon microscopy. Opt. Express 23, 13833–13847 (2015).

16. Bumstead, J. R., Park, J. J., Rosen, I. A., Kraft, A. W., Wright, P. W., Reisman, M. D., Côté, D. C., and Culver, J. P., Designing a large field-of-view two-photon microscope using optical invariant analysis. Neurophoton. 5, 025001 (2018).

17. Terada, S. I., Kobayashi, K., Ohkura, M., Nakai, J., and Matsuzaki, M., Super-wide-field two-photon imaging with a micro-optical device moving in post-objective space. Nature Commun. 9, 3550 (2018).

18. Stirman, J. N., Smith, I. T., Kudenov, M. W., and Smith, S. L., Wide field-of-view, multi-region, two-photon imaging of neuronal activity in the mammalian brain. Nature Biotechnology 34, 857–862 (2016).

19. Balu, M., Mikami, H., Hou, J., Potma, E. O., and Tromberg, B. J., Rapid mesoscale multiphoton microscopy of human skin. Biomedical Opt. Express 7, 4375–4387 (2016).

20. Sofroniew, N. J., Flickinger, D., King, J., and Svoboda, K., A large field of view two-photon mesoscope with subcellular resolution for in vivo imaging. eLife 5, e14472 (2016).

21. Prevedel, R., Verhoef, A. J., Pernĺa, A. J., Weisenburger, S., Huang, B. S., Nöbauer, T., Fernández, A., Delcour, J. E., Golshani, P., Baltuska, A., and Vaziri, A., Fast volumetric calcium imaging across multiple cortical layers using sculpted light. Nature Methods 13, 1021–1028 (2016).

22. Weisenburger, S., Tejera, F., Demas, J., Chen, B., Manley, J., Sparks, F. T., Traub, F. M., Daigle, T., Zeng, H., Losonczy, A., and Vaziri, A., Volumetric Ca2+ Imaging in the Mouse Brain Using Hybrid Multiplexed Sculpted Light Microscopy. Cell 177, 1050–1066 (2019).

23. Zipfel, W. R., Williams, R. M., and Webb, W. W., Nonlinear magic: multiphoton microscopy in the biosciences. Nature biotechnology 21,1369–1377 (2003).

24. Sheppard, C., and Gu, M., Image formation in two-photon fluorescence microscopy. Optik 86,104–106 (1990).

25. Jaepel, J., Hübener, M., Bonhoeffer, T., and Rose, T., Lateral geniculate neurons projecting to primary visual cortex show ocular dominance plasticity in adult mice. Nature Neuroscience 20, 1708–1714 (2017).

26. Chun, W., Do, D., and Gweon, D. G.,Design and demonstration of multimodal optical scanning microscopy for confocal and two-photon imaging. Review of Scientific Instruments 84, 013701 (2013).

27. Kong, L., Tang, J., Little, J. P., Yu, Y., Lämmermann, T., Lin, C. P., Germain, R. N., and Cui, M., Continuous volumetric imaging via an optical phase-locked ultrasound lens. Nature Methods 12, 759–762 (2015).

28. Gil, H. H., Golgher, L., Israel, S., Kain, D., Cheshnovsky, O., Parnas, M., and Blinder, P., PySight: plug and play photon counting for fast continuous volumetric intravital microscopy. Optica 5, 1104–1112 (2018).

29. Li, B., Wu, C., Wang, M., Charan, K., and Xu, C., An adaptive excitation source for high-speed multiphoton microscopy. Nature Methods 17, 163-166 (2020).

30. Xiao, S., and Mertz, J., Contrast improvement in two-photon microscopy with instantaneous differential aberration imaging. Biomed. Opt. Express 10, 2467–2477 (2019).

31. Charan, K., Li, B., Wang, M., Lin, C. P., and Xu, C., Fiber-based tunable repetition rate source for deep tissue two-photon fluorescence microscopy. Biomed. Opt. Express 9, 2304–2311 (2018).

32. Chung, K., and Deisseroth, K. (2013). CLARITY for mapping the nervous system. Nat Methods 10, 508–513.

